# Molecular diffusion and binding kinetics are sufficient to explain condensate-like nuclear clustering

**DOI:** 10.64898/2026.06.04.730171

**Authors:** Manya Kapoor, Mustafa Mir

## Abstract

Many key transcriptional regulators form clusters in cell nuclei that are proposed to promote transcription by concentrating reactants. These clusters exhibit emergent biophysical properties consistent with biomolecular condensates such as high local densities, spherical morphologies, fusion, and rapid molecular exchange kinetics. Whether these properties are specific hallmarks of condensation or can arise from simpler mechanisms is unclear. Here we developed SPARK (Simulation of Protein Accumulation from Reaction Kinetics), a computational framework parameterized by genomic, and molecular diffusion and binding kinetics data, to simulate the spatial distributions of chromatin-binding proteins as they would appear under an optical microscope. These simulations recapitulate key reported features of Brd4, Nanog, Sox2, and Bicoid clusters, without including any higher-order protein-protein interactions or phase separation. We demonstrate that commonly used signatures of condensates instead emerge directly from molecular scale binding and diffusion kinetics. Through systematic variation of nuclear concentrations and enhancer binding-site composition, we show that distinct combinations of molecular kinetics can result in similar spatial distributions of transcription factors. These analyses establish that imaging-based features of protein clusters cannot uniquely define the underlying physical mechanism of their formation. We therefore conclude that condensation is not required to drive transcription factor occupancy and that protein-protein interactions off chromatin are not required to explain cluster formation. Our results instead imply that mechanisms other than clustering are sufficient for target-search acceleration and robust occupancy. SPARK thus provides a quantitative null model for identifying which molecular mechanisms are necessary to explain experimental observations of nuclear protein clustering.

## INTRODUCTION

Transcription factors (TFs) regulate gene activity by binding to regulatory genomic regions known as enhancers and promoters. Although bursts of transcriptional activity at genes can last for minutes to hours, live-cell single-molecule tracking experiments have shown that TFs generally bind chromatin transiently, on the order of just tens of seconds^1–5^. Despite these short binding times, many TFs have been shown to form clusters, often referred to as condensates or hubs, that can persist over much longer time-scales^3,4,6,7^. Such cluster formation has also been observed for RNA Polymerase II (RNAPII) and transcriptional co-activators, suggesting that such spatial organization of regulatory proteins is a widespread feature of transcriptional regulation^8–11^. These observations have challenged canonical models of transcription factor function in which stable binding followed by hierarchical recruitment of co-factors is required for regulating transcriptional activity. Instead, clusters are proposed to be higher-order regulatory structures that promote sustained transcription factor occupancy by concentrating factors around genes and driving continuous and rapid exchange^3,4,9,10,12,13^. Understanding how these local nuclear microenvironments form and persist is therefore critical to establishing the molecular mechanisms underlying transcriptional regulation.

A proposed mechanism for driving clustering is biomolecular condensation through phase separation driven by multivalent interactions between TFs and their co-factors^9,10,14,15^. Studies on Mediator, RNAPII, and Brd4 have led to a model in which condensates concentrate factors and drive transcriptional activity^9,10,12,13,15–18^. Condensates are often characterized using imaging based criteria such as spherical shape, fusion events, and rapid fluorescence recovery after photobleaching^9,10,19,20^. However, similar clustering behavior can emerge through other mechanisms^14,21^. For example, weak and transient homo- and heterotypic interactions can lead to local enrichment without phase separation^11^. Molecules can also appear to accumulate in large domains due to a spatial clustering of their substrates^22,23^. Recent work from our lab and others has also shown that the appearance of condensate-like bodies may be explained more simply by molecular diffusion and binding kinetics and limitations of microscopy^6,8,22,24^. A quantitative null model is therefore needed to distinguish the properties of nuclear clusters which require higher-order molecular interactions to explain, from those that do not. A model containing only the simplest processes required to produce an observation provides an important baseline to establish whether additional physical processes are required.

Here we developed a computational framework dubbed SPARK (Simulation of Protein Accumulation from Reaction Kinetics) to simulate the spatial distribution of TFs from molecular diffusion and binding kinetics. SPARK generates both molecular trajectories and corresponding microscopy images, allowing for direct comparisons between molecular scale kinetics and experimentally observable data. SPARK does not simulate multi-valent protein-protein interactions or phase separation allowing us to assess which emergent condensate properties can be predicted from diffusion and binding kinetics alone. We applied SPARK to reproduce reported condensate formation across a range of transcriptional regulators and biological systems. Additionally, we simulated FRAP (Fluorescence Recovery After Photobleaching), apparent cluster fusion, and the effects of binding site properties on TF occupancy. Together, our analyses establish a quantitative null model for interpreting nuclear clustering or condensate formation.

## RESULTS

### SPARK enables modulation of kinetic and microscopy parameters

SPARK is packaged as a user-friendly interactive tool (Fig. S1). The software allows users to specify a range of experimentally-measurable molecular kinetic parameters which are used to generate molecular trajectories. Each molecule undergoes Brownian diffusion until it encounters a binding site, to which it can bind with a user-defined probability k^on^. Users can specify the number, clustering, and strength of binding sites, as well as introduce synthetic binding site arrays. The binding sites diffuse via fractional Brownian motion to recapitulate constrained polymer motion, and upon successful binding, the molecule diffuses along with the binding site. The duration of each binding event is drawn from a user-defined residence time distribution after which the molecule unbinds and resumes free diffusion (Fig. 1A).

**Figure 1.**
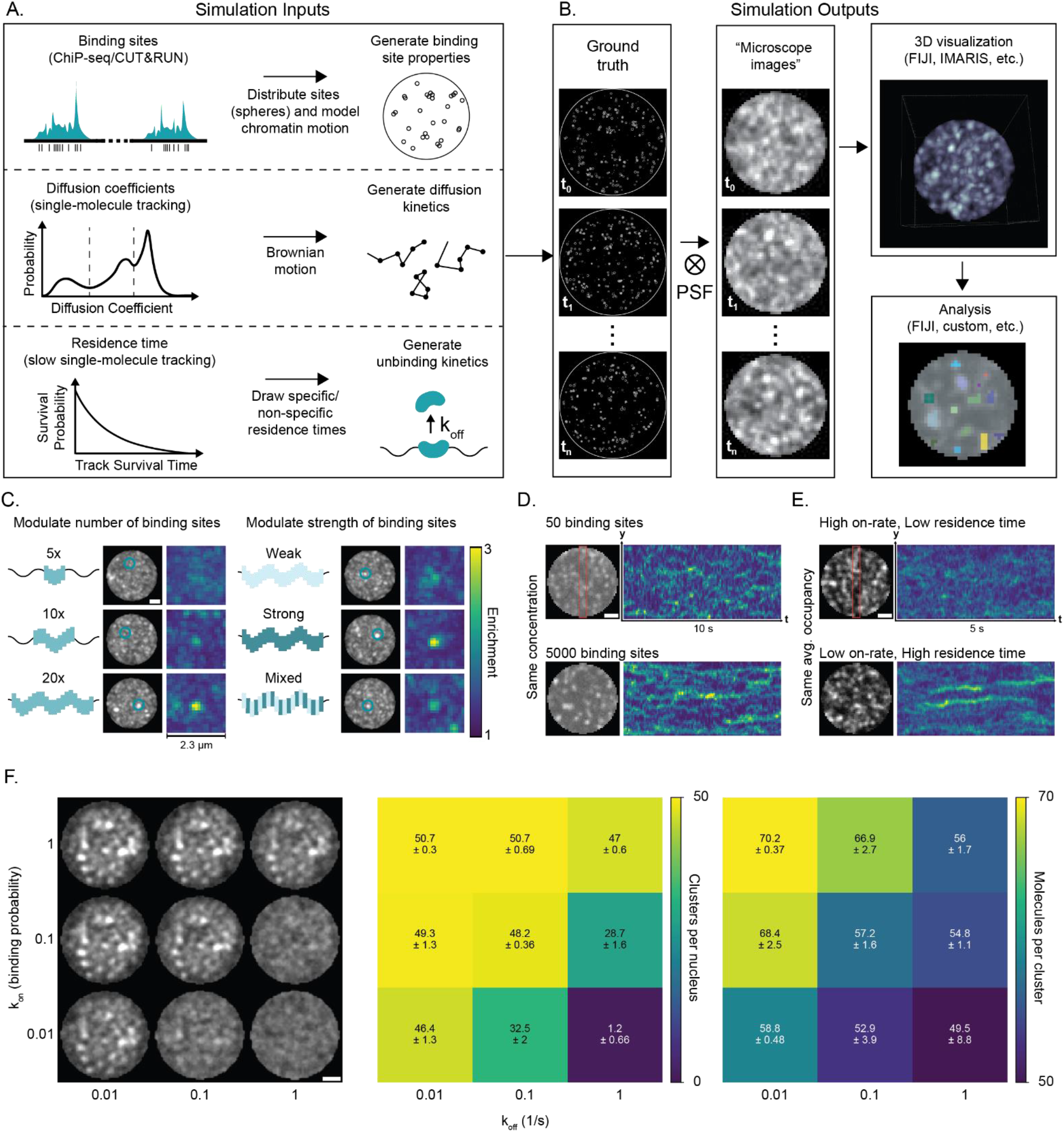
SPARK enables modulation of kinetic and microscopy parameters. **(A)** Simulation inputs. Users input binding site characteristics, molecular diffusion coefficients, and residence times. **(B)** Simulation outputs. The simulator generates molecular positions which are convolved with a user-defined point spread function (PSF) to obtain z-slices of “microscope images” at each time point. These can be visualized in any image analysis software and further analysed. **(C)** Simulating binding sites with varying numbers/strengths. Left: Binding sites schematic. Middle: Simulated nucleus. Blue circle indicates binding site array position. Right: Enrichment of TF over nuclear background, centered at the binding array. **(D)** Nucleus image (left) and kymographs over time (right) in the max-projected red area showing the effects of changing the number of binding sites in the nucleus. Scale bar is 1 μm. **(E)** Nucleus image (left) and kymographs (right) in the maximum intensity projected red area showing the effects of changing the TF on-rate and residence times. Scale bar is 1 μm. **(F)** Simulated nuclei (left) and heatmaps of number of clusters (middle) and number of molecules per cluster (right) for a range of k_on_ and k_off_ values. Scale bar is 1 μm. Numbers show mean and standard deviation across three replicates.

SPARK generates two visual outputs: “ground truth” images of molecular positions and binding sites, and simulated microscope images (SI Methods). The latter is produced by convolving the molecular position data with the microscope’s point spread function (PSF) and mapping the results to discrete pixel intensities, mimicking a digital camera sensor. This approach is used to generate simulated time-lapse volumetric imaging data, taking into account variables such as z-slice spacing, time between frames, and exposure time. The ground truth and simulated microscope images can be viewed on a browser and downloaded as multi-page TIFFs for further analysis (Fig. 1B). Simulations can be seeded at two levels: identical molecular kinetics, which allows modulation of microscopy visualization parameters, and identical binding site kinetics, which allows modulation of molecular kinetics alongside visualization parameters (SI Methods). As a commonly used experiment to assess the biophysical properties of cluster and biomolecular condensates is Fluorescence Recovery After Photobleaching (FRAP), SPARK includes a module to perform computational FRAP experiments on regions of interest within simulated images. SPARK can also output molecular positions for reconstruction of super-resolution microscopy data.

We first used SPARK to design enhancers with varying affinities and observe the resulting TF clustering. Consistent with previous observations^6^, we find that modulating the number and strength of binding sites at a single locus affects the enrichment of the TF at the locus over the nuclear background, leading to the appearance of a bright, stable cluster (Fig. 1C). We then systematically modulated individual nuclear parameters to gain intuition on how they alter the appearance of clusters. Increasing the total number of binding sites globally in the nucleus increases the visibility and persistence of clusters (Fig. 1D). We then modulated molecular kinetic parameters and found that similar clustering can be achieved by modulating either the on-rate (here defined as the probability that a molecule in the vicinity of a binding site will bind) or the residence time of the molecule (Fig. 1E). Across a broad range of association probabilities and residence times, cluster numbers remain similar despite substantial differences in molecular occupancy. However, combining low association probabilities with short residence times markedly reduces detectable clustering (Fig. 1F). Importantly, altering microscope parameters (exposure time, resolution, etc.) has a profound impact on how clusters appear with identical underlying molecular kinetics. For example lower spatial or temporal resolution leads to the appearance of larger and more spherical clusters (Fig. S2). Thus, apparent cluster properties depend on a range of nuclear, molecular kinetic, and microscopy parameters.

### Concentration dependent clustering and condensate properties are recapitulated by molecular kinetics

To test the effect of increasing concentration on clustering behavior, we simulated nuclei with identical binding site kinetics and molecular dynamics along a concentration gradient ranging from 200-550 nM (Fig. 2A), which is typical of many nuclear TFs, such as Dorsal in the *Drosophila* embryo^6,25^. Using image segmentation to characterize clusters^6^, we found that the number of clusters is relatively constant across the gradient (Fig. 2B, left). However, filtering for high-confidence clusters (enrichment over nuclear background > 1.15) resulted in a drop in cluster number at lower concentrations (Fig. 2B, right). Using this threshold, we found that the average cluster intensity increases as a function of concentration, similar to experimental findings for the TF Dorsal in *Drosophila* embryos^6,26^, where cluster intensity scaled with nuclear intensity (Fig. 2C). Finally, we counted the number of molecules per cluster and saw a wide distribution, which increased with increasing concentration (Fig. 2D). The majority of clusters had fewer than 100 molecules per cluster at all concentrations and were distinguishable from nuclear background, indicating that high density phase transitions are not necessary to produce visible clusters.

**Figure 2.**
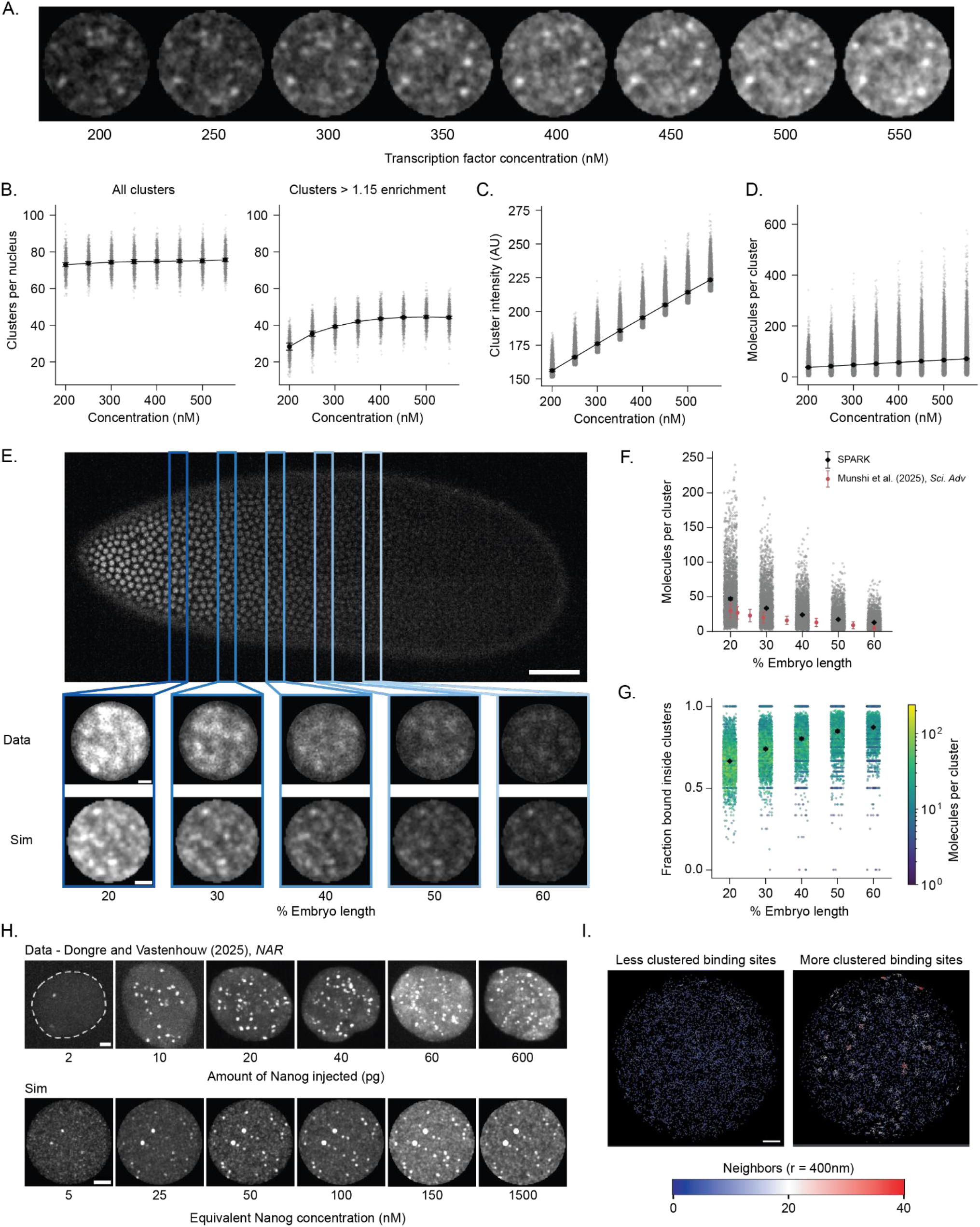
Dependence of clustering on nuclear concentrations. **(A)** Sample images (single frame, single z-slice) of nuclei with identical binding site distributions and molecular kinetics but different concentrations. Each nucleus is 5 μm in diameter. **(B)** Segmented clusters at each concentration with (right) and without (left) enrichment thresholding. **(C)** Cluster intensity vs. nuclear concentration. **(D)** Molecules per segmented cluster vs. nuclear concentration. Data from 5 simulations for each concentration are shown for (B-D), and error bars indicate SD across replicates. **(E)** Bicoid gradient. Top: Maximum intensity projection image of Bicoid-GFP in a Drosophila embryo in nc14. Scale bar is 50 μm. Bottom: Data vs. simulated images of individual nuclei (single frame, single z-slice). Scale bars are 1 μm. **(F)** Number of Bicoid molecules per cluster along the Bicoid gradient for simulated data and previously estimated data^29^. Error bars indicate SD (n=5 replicates for simulated data). **(G)** Fraction of bound molecules (both specific and non-specific) in each segmented cluster. The color of the points indicate the number of molecules per cluster. Error bars indicate SD across 5 replicates for simulated data. **(H)** Data vs. simulations for 512-cell stage zebrafish embryo nuclei injected with varying Nanog concentrations. Images are maximum intensity projections in z and contrast adjusted to highlight clustering. Scale bars are 2 μm. **(I)** Density rendering of bound Sox2 binding in ES cells with low (left) and high (right) clustering of binding sites. Colorbar indicates the number of bound Sox2 molecules within a 400 nm radius sphere around each molecule. Scale bar is 1 μm.

We next tested if SPARK can reproduce experimentally measured concentration dependent clustering. We first examined Bicoid, a morphogen TF that forms a concentration gradient along the anterior-posterior axis of the *Drosophila* embryo, and is known to cluster at both high and low nuclear concentrations across a range of 5-1000 nM, as estimated by recent work^3,4,27–29^. Parameterizing SPARK using published data^3,4,28,29^, we simulated nuclei at various points along the Bicoid concentration gradient and found that SPARK recapitulates clustering across all concentrations (Fig. 2E). Consistent with experimental observations and parameter sweeps, we found that the number of clusters decreases at lower concentrations, suggesting that some clusters of binding sites are not saturated (Fig. S3A). Using SPARK’s ground truth molecular position data, we counted the molecules within each cluster, and found that the number of molecules within simulated clusters shows a linear relationship with concentration, consistent with previous estimations^29^ (Fig. 2F). Interestingly, the fraction of bound molecules per cluster increases as concentration decreases, suggesting that full occupancy for most clusters can be achieved at low concentrations, and the additional molecules are free molecules in the nuclear background contributing to cluster intensity (Fig. 2G). This is consistent with single-molecule data showing that the overall bound fraction of Bicoid in posterior nuclei is higher than in anterior nuclei^4^. Additionally, we show that while other estimates of Bicoid nuclear concentration lead to different numbers of clusters per nucleus, or molecules per cluster, the observation of clustering remains consistent (Fig. S3B-C).

We next used SPARK to simulate Nanog in zebrafish embryos. Nanog is a transcription factor implicated in maintaining pluripotency in zebrafish embryos, and in human and mouse embryonic stem cells. It forms dynamic clusters at transcriptional bodies, where it colocalizes with RNAPII and elsewhere in the nucleus. In previous work, Dongre et al. injected varying concentrations of Nanog mRNA into one-cell stage zebrafish embryos, and imaged them at the 512-cell stage^30^. They concluded that the clustering of Nanog is dependent on both its DNA-binding domain and at least one Intrinsically Disordered Region (IDR). We parameterized SPARK using experimentally measured estimates of nuclear size^30–32^, number of binding sites^31,33–35^, and Nanog kinetics^2,36^, and a calculation of injection and translation efficiency, and simulated 512-cell stage nuclei injected with the calculated Nanog concentrations. We see that SPARK reproduces strikingly similar images to those produced by confocal microscopy (Fig. 2H). Our results suggest that the reported IDR dependence on Nanog’s clustering may not be driven by multivalent interactions which lead to clustering of unbound molecules, but rather IDR mediated tuning of binding kinetics. This is in line with the observation that Nanog clusters at the miR-430 locus are seeded by the locus^31^.

Next, to test SPARK’s ability to simulate super-resolution microscopy data we examined the clustering of Sox2, a key pluripotency regulator known to have clustered binding sites^2^. Using the ground truth molecule positions that SPARK outputs, we reconstructed a 3D rendering of bound Sox2 molecules in ES cells and colored each molecule by its number of neighbors. Consistent with published data we find that Sox2 clusters only become visible upon clustering of its binding sites^37^ (Fig. 2I, Movie S1). Together these three case studies show that the appearance and characteristics of TF clusters can be recapitulated through diffusion, binding kinetics, and the properties of binding sites.

### Molecular kinetics simulations reproduce key quantitative and qualitative features of Brd4 transcriptional condensates

We next simulated the transcriptional coactivator Brd4, which has been described in the literature to form phase-separated condensates^10,38–40^. Brd4 is a mammalian bromodomain and extra-terminal (BET) protein that binds acetylated histones and is implicated in clustering transcriptional machinery at active genes^10,38^. Single molecule tracking in mouse ESCs shows that Brd4 is present in bound, free, and intermediate diffusion populations within the nucleus^41^. To simulate Brd4, we used a three-state molecular kinetics model and swept the exchange rates between the states (Fig. 3A). We ran 243 simulations and calculated the resulting bound, free and intermediate fractions for each. We compared these fractions to the experimentally measured fractions for Brd4^41^, and identified the parameter that most closely matched the experimental data (Fig. 3B-D). Using this parameter set we simulated a time-series of three-dimensional Brd4 images. To test whether we could recapitulate reported recovery kinetics of condensates, we performed a simulated FRAP experiment on a Brd4 cluster at similar timescales as those reported for Brd4 condensates. We found that our simulated recovery kinetics closely match experimentally reported ones^10,20^ (Fig. 3E-F, Movie S2). Moreover, in our simulated data, we were able to readily observe local accumulations that resemble liquid-like droplets, and that undergo fusion events similar to experimental observations of condensate fusion^10,15^ (Fig. 3G, Movie S3). Importantly, in the simulations all fusion events are due to chromatin motion not droplet fusion. The integrated intensity of the “fused” cluster is approximately the sum of the two individual pre-fusion clusters, an observation commonly attributed to condensate coalescence (Fig. 3H). These results demonstrate that FRAP recovery kinetics and apparent fusion, two key criteria commonly used to classify clusters as condensates, can arise from molecular binding kinetics and chromatin motion alone.

**Figure 3.**
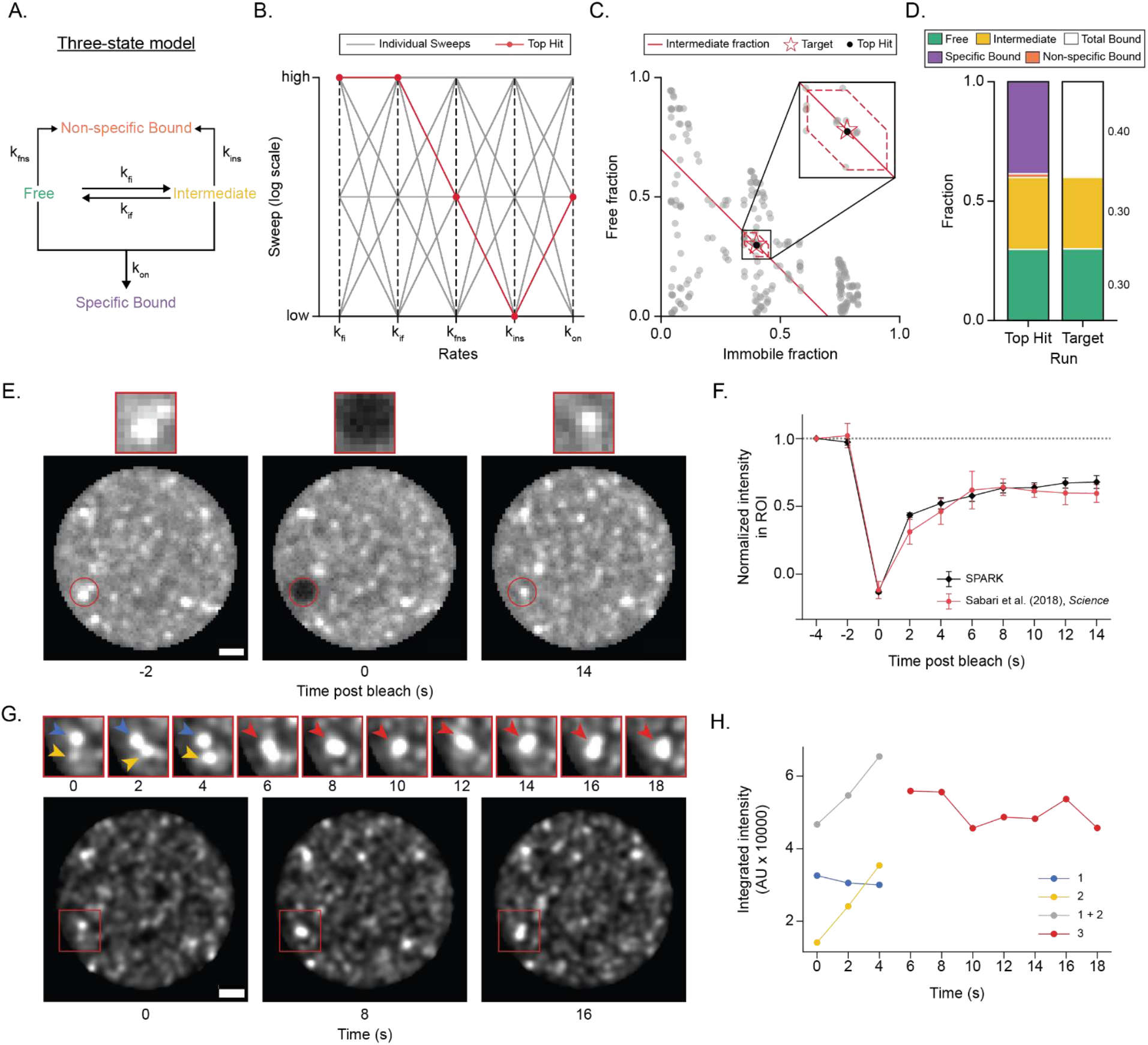
Clusters arising from Brd4 molecular kinetics simulations exhibit properties attributed to condensates. **(A)** Three-state model of kinetic populations. **(B)** Landscape of parameter sweeps (grey lines). 3^5^ = 243 total simulations were run to cover each combination of the five rates. **(C)** Bound, free, and intermediate fractions resulting from each sweep (grey dot). The red star indicates the target to match (Xie et al. 2022)^41^. Hexagon marks the region where fractions fall within 0.05 of the target. **(D)** Bound, free, and intermediate fraction of the run closest to the target. **(E)** Sample images for a simulated FRAP experiment. Red circle depicts a 1 μm diameter photobleached region in which intensity is measured. Insets show zoomed-in views of the red circles. Scale bar is 1 μm. **(F)** Average FRAP intensity traces from simulations (n=5) and experimental (n=9) data (Sabari et al. 2018)^10^. Error bars show SEM for both simulated and experimental data. **(G)** Sample images showing clusters exhibiting fusion. Insets show zoomed-in views. Scale bar is 1 μm. **(H)** Cluster intensities of the individual and fused clusters.

### Different enhancer compositions can result in similar TF occupancy and clustering

Across various organisms, cell fate specification and body plan development occur through the establishment of spatial and temporal concentration gradients by transcription factors known as morphogens^42,43^. It has been suggested that enhancers comprising weak binding sites can achieve specificity and, when sufficiently clustered, can produce equivalent levels of gene expression as strong binding sites ^44–48^. SPARK allows modulation of concentration and enhancer strength, clustering, and cooperativity, and can be used to test how these parameters modulate TF occupancy and the appearance of clustering.

We first used SPARK to test whether strong binding sites (sites with high residence times), weak binding sites (sites with low residence times), or combinations of both can result in varying points of activation along spatial or temporal concentration gradients. We did not vary association rates between weak and strong binding sites. We modeled a cluster of six binding sites and set an activation threshold at 1/3rd sites occupied. We found that stronger sites reached this threshold at lower concentrations and thus cannot substantially increase their occupancy at higher concentrations. Clusters of weaker sites are not fully occupied at low concentrations and thus have more capacity to respond to concentration changes. Mutating a weak site to a stronger one shifted its activation threshold towards a lower concentration (Fig. 4A). This result is in line with the observation of strong and weak sites in enhancers that respond to morphogens like Bicoid in the *Drosophila* embryo at different positions along its concentration gradient^42^.

**Figure 4.**
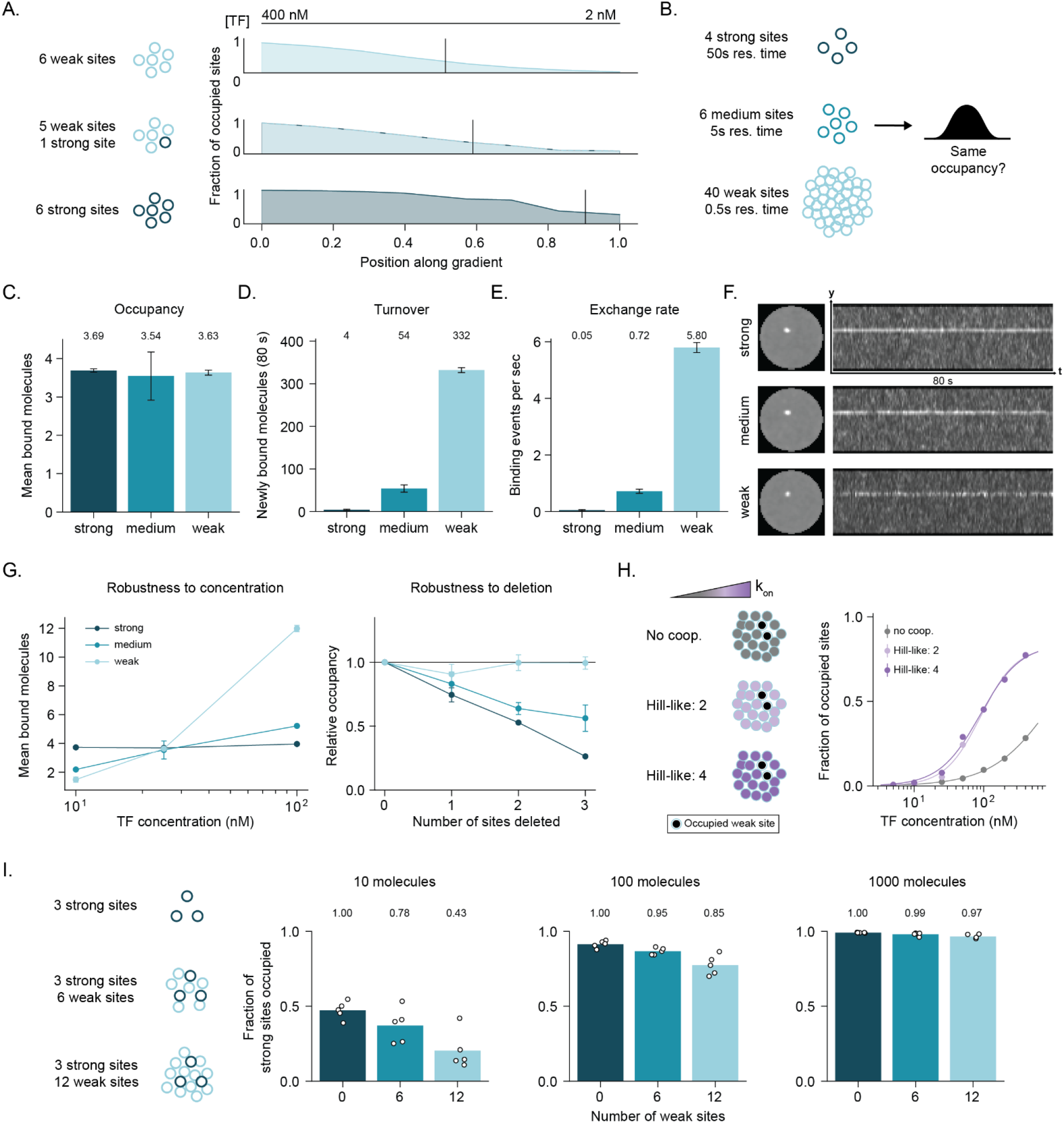
Clusters of weak binding sites have tunable occupancy. **(A)** Fraction of occupied sites in a cluster of six strong, weak, or mixed binding sites as a function of TF concentration. Vertical line indicates concentration at which 1/3rd of sites are occupied on average. **(B)** Schematic for tested hypothesis in (C-G). **(C)** Average occupancy over 80s for each cluster of binding sites in (B), from two independently simulated replicates. Error bars indicate SD. **(D)** Molecular turnover over 80s for each cluster of binding sites in (B), from two independently simulated replicates. Numbers above the bars indicate the value. Error bars indicate SD. **(E)** Molecular exchange rate for each cluster of binding sites in (B), from two independently simulated replicates. Error bars indicate SD. **(F)** Maximum intensity projections over time (left) and kymographs (right) showing the cluster arising from the sites in (B). A three-slice maximum projection is done in z to capture the cluster of binding sites. **(G)** Left: Average occupancy for each cluster in (B) as a function of nuclear TF concentration. Right: Fractional occupancy of each cluster in (B) as a function of number of sites deleted from the cluster. Error bars indicate SD. **(H)** Schematic (left) and plot showing average occupancy as a function of nuclear TF concentration for clusters of weak binding sites with varying levels of cooperativity. **(I)** Schematic (left) and plots showing average occupancy at strong binding sites with varying numbers of weak binding sites in the same cluster at low (left), medium (middle) and high (right) TF concentrations. Dots indicate individual simulated replicates (n=5). Error bars indicate SD. Numbers above the bars indicate the value.

Since strong sites can achieve full occupancy at lower concentrations, we wondered if having many weak sites could provide a greater opportunity for tuning occupancy, and may explain why clusters of weaker redundant binding sites may be favoured over fewer, stronger binding sites in an evolutionary context^48^. We used SPARK to test whether similar high target occupancy, and resultant apparent clustering, could result from either a small number of strong binding sites or a large number of weak binding sites (Fig. 4B). After sweeping various combinations and calculating occupancy over time, we found that similar levels of occupancy could be obtained with four strong binding sites, six medium-strength binding sites, or forty weak sites, with their residence times differing by an order of magnitude each (Fig. 4B-C). As expected, the weak sites had a high turnover of over 300 molecules in 80 seconds, while the strong sites had a turnover of just four molecules over the same time window (Fig. 4D-E). While visually forming similar clusters, analysis of kymographs revealed that while instantaneous occupancy is at times higher at the large number of weak sites, the large amount of turnover led to a similar time averaged occupancy as the fewer strong sites (Fig. 4F). Interestingly, modulating the nuclear concentration revealed that weaker sites are more sensitive to changes in concentration, with average occupancy changing by a factor of eight over an order of magnitude change in concentration, while the stronger sites showed the same average occupancy across this concentration range (Fig. 4G). Conversely, the weaker sites were more robust to site deletions (Fig. 4G). This could explain why low-affinity binding sites are favored over stronger ones in evolutionary contexts – they respond to morphogen gradients and their average occupancy is more resistant to site mutations^44,47^.

We next tested whether small levels of homotypic cooperative binding (defined in SPARK as the binding of one molecule increasing the on-rate of molecules binding to the neighboring vacant sites in the cluster of binding sites) could lead to switch-like transitions in TF occupancy at a locus of weak sites. We found that incorporating a small increase in cooperativity (Hill-like coefficients of 2 and 4) makes a cluster of weak sites more sensitive, leading to a near switch-like occupancy transition at a critical concentration (Fig. 4H). Thus, cooperativity is a mechanism by which occupancy can be modulated at weak binding sites.

Finally, we tested whether the presence of weak sites can increase occupancy of stronger sites in the same cluster of binding sites as has been proposed in previous work^48,49^. To test this, we simulated an enhancer with three strong sites and either zero, six, or 12 weak binding sites in their vicinity (Fig. 4I). However, the presence of weak sites did not enhance average occupancy at the stronger ones, but rather reduced it at low concentrations (Fig. 4I). Since SPARK only simulates 3D diffusion, the weak sites shield the strong ones – once a molecule unbinds, it is free to explore the entire nucleus rather than being funneled towards the stronger sites. As we do not incorporate theories of facilitated diffusion such as 1D sliding and hopping, or other methods of local confinement these results may not reflect mechanisms which lead to the occupancy of strong sites surrounded by weak sites in vivo^50–53^.

## DISCUSSION

Our results demonstrate that several imaging-based features commonly associated with transcriptional condensates can emerge from molecular diffusion and binding kinetics. Since SPARK does not account for protein-protein interactions, it cannot directly rule out higher-order processes of cluster formation. Instead, it asks whether observed clusters can be explained without needing to invoke higher-order mechanisms for cluster formation. Using SPARK, we reproduced key features of nuclear condensates formed by transcriptional regulators across different biological systems, including local enrichment, cluster lifetimes, apparent fusion events, and FRAP. These results highlight that the properties of a bright cluster in microscopy data and the physical mechanism responsible for its formation must be carefully distinguished. For example the apparent fusion or merging of two fluorescent clusters does not necessarily reflect droplet coalescence, and rapid FRAP recovery can arise from independently binding molecules. Although such observations may be consistent with biomolecular condensation, they are not specific to it^11,14,21,22^.

Our simulations show that the spatial distributions and properties of binding sites, coupled with molecular kinetics, can lead to variation in the emergent properties of nuclear clusters. Increasing binding site number, residence times, and association rates, can all lead to clustering, but with very different underlying effects on molecular turnover and sensitivity to nuclear concentration. For example, we show that similar time averaged occupancy can arise from a few strong sites or a large number of weak sites. Furthermore an enhancer containing a large number of weak sites is both more sensitive to changes in nuclear concentration and robust to perturbations to binding site number. This is consistent with observations of developmental enhancers containing clusters of low affinity binding sites across organisms^44,46–48^. These findings also suggest that enhancer composition tunes the dynamics and robustness of TF occupancy consistent with our recent experimental observations of Dorsal in *Drosophila* embryos^6^. More broadly, stable local enrichment of a TF can be achieved through the continuous exchange of molecules without requiring long-lived binding of individual TFs or condensate formation.

Our analysis also raises a broader question on how TFs locate their genomic targets. Current models often invoke facilitated diffusion in which DNA-binding proteins can accelerate their target search by combining three-dimensional diffusion with one-dimensional sliding or other forms of local exploration^2,49–53^. In live eukaryotic cells, many variables are at play, including fluctuations in chromatin accessibility, non-specific interactions, and protein-protein interactions that may help locally retain molecules. Although SPARK includes Brownian motion, non-specific binding, and a probabilistic mode of target “capture”, association probabilities are defined and not dependent on any molecular mechanisms. As such, SPARK can reproduce experimentally observed clustering but cannot be used to directly conclude whether target search is reaction or diffusion-limited. However, it is important to distinguish the time required to encounter a target versus the time required to productively bind to it. As such an important implication of SPARK is that TFs may rely on additional mechanisms that help facilitate effective target recognition and reduce non-specific interactions^54^. These mechanisms are independent of the free diffusion coefficient or the necessity to form a condensate.

Intrinsically disordered regions (IDRs) are ideal candidates for protein domains that contribute to TF target localization and binding. Although IDRs are frequently implicated in phase separation and condensate formation due to their propensity for multivalent interactions, their contributions to TF function extend beyond clustering. For example, IDRs regulate genomic binding specificity^55,56,57^, promote transient confinement^53^, facilitate target search through transient interactions with chromatin^58,59^, target scanning^60^, and tuning association and dissociation kinetics at specific vs non-specific sites^52^. These studies demonstrate that IDRs can engage in molecular interactions that increase the probability of productive target occupancy without phase separation. Although SPARK does not directly simulate IDR interactions, the molecular kinetics parameters used to constrain the simulations may capture these effects as they are derived from experimental data. Similarly, the association probabilities used in SPARK may reflect IDR mediated local-confinement or recruitment. The fact that SPARK reproduces nuclear clustering without encoding such interactions does not imply that they do not exist or are not important. Rather, these molecular interactions likely determine the kinetic parameters that govern local TF occupancy without the need for condensation.

In conclusion, SPARK demonstrates that chromatin binding and diffusion kinetics on their own are able to recapitulate the appearance and biophysical properties of condensate-like bodies formed by transcription factors. We conclude that clusters that might appear to be liquid-like condensates in fluorescence microscopy data can arise from molecular binding kinetics. SPARK will allow others to similarly assess whether the clusters they observe can be explained by chromatin binding and diffusion kinetics, rather than higher-order phenomena like condensate formation.

## METHODS

### Simulator overview

SPARK is a browser-based tool implemented in Streamlit/Python, used to generate ground truth and “microscope-like” images from molecular data. The simulator consists of an interactive mode, which allows users to simulate and visualize results from a single set of parameters in the browser, and a batch mode where users can upload a CSV specifying a range of simulation parameters, the results of which can be downloaded. A single physics engine drives both modes. Additionally, a command-line implementation of the same engine enables running simulations on high-performance computing cores (see SI Methods).

#### Geometry and molecule count

The nucleus and transcription factor (TF) molecules are modelled as spheres. The number of TF molecules is set from the user-supplied bulk concentration and the nuclear volume:

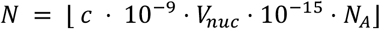

with *c* in nM, *V*_*nuc*_ in µm^3^ and *N*_*A*_ = 6.022 × 10^23^, or given directly. All distances internal to the engine are tracked in micrometres, while nanometres are used for image-space parameters (PSF, voxel size, slice spacing).

#### Time stepping and exposure averaging

Three timescales coexist:

- **Physics step** *Δt*: the timestep at which positions, states, and binding events are updated. By default this is set at the exposure time for single molecule tracking experiments (10 ms).
- **Exposure time** *T*_*exp*_: a contiguous block of *T*_*exp*_/*Δt* physics steps representing camera integration. At every step within the window, a snapshot of all molecule positions, states, site positions, and site occupancies is recorded.
- **Display interval** *T*_*disp*_ : the period at which images are generated. Each display interval begins with an exposure window. The remaining physics steps are “dark” steps that advance the system without generating an image.

For each output frame, molecule positions are compressed to a single per-molecule centroid by averaging across the exposure window. Molecule states (see below) are compressed by mode (most frequent state during the window); site positions are averaged; site occupancy is the logical OR across the window (a site is marked occupied if it was occupied at any point during exposure). This produces a faithful integration of motion and switching during the camera exposure rather than a single instantaneous snapshot.

#### Molecule states and dynamics

##### State space

The simulator supports two state schemes:

- **3-state**: molecules occupy one of: Free, Intermediate, and Bound states. The Bound state consists of both specific binding to binding sites and non-specific (NS) binding.
- **2-state**: the Intermediate state is removed; only Free and Bound states (both specific and non-specific) exist.

##### Initial populations

The user supplies the initial percentages of mobile molecules in Free, Intermediate, and Non-Specifically bound states. Initial positions are drawn uniformly inside the nucleus and states are sampled from the supplied distribution.

##### Per-state diffusion

Each molecule (with the exception of specifically bound molecules) is assigned a diffusion coefficient *D* drawn from a Gaussian whose mean and standard deviation depend on its state. *D* is drawn at initialization and again whenever the molecule’s state changes. Negative draws are clipped to zero. Specifically bound molecules are pinned to their cognate site for the duration of binding and move along with the site.

##### First-order state transitions

At every *Δt*, mobile molecules undergo Markov transitions at user-specified probabilities. NS-bound molecules return to the mobile pool with probability *Δt*/*τ*_*NS*_ where *τ*_*NS*_ is the user-specified mean non-specific residence time (1/*τ*_*NS*_ = *k*_*off*_ or the non-specific off-rate). After unbinding, molecules get assigned to the Free or Intermediate state with equal probability (in 2-state mode the destination is always Free). Newly transitioned molecules are immediately re-assigned a state-appropriate diffusion coefficient.

##### Specific binding

At each *Δt*, the positions of mobile molecules are queried against a *k*-*d* tree of binding-site coordinates (rebuilt only when sites have moved). Any mobile molecule within the user-set capture radius of the closest site can bind with probability *p*_*on*_ if the site is free. When cooperative binding is enabled, this probability becomes *p*_*on*_ ⋅ [1 + (*αn*)^h^] clipped to 1, where *n* is the number of already-bound sites in that cluster, *α* is the cooperativity strength, and h is the Hill coefficient. Multiple molecules competing for the same site are resolved by a random shuffle and uniqueness pass such that each site can only be occupied by one molecule. On binding, the molecule is locked to the site position, the cluster’s occupancy counter increments, and an unbind time is drawn:

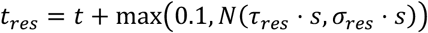

where *s* is the site strength multiplier and *τ*_*res*_, *σ*_*res*_ are user-set residence-time mean and standard deviation. Bound molecules are returned to the mobile pool (Free or Intermediate state) with equal probability.

##### Diffusive step and boundary

Mobile molecules are propagated by an isotropic Gaussian step with per-axis standard deviation 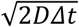. Any mobile molecule whose updated position lies outside the nuclear sphere is projected radially back onto the surface.

#### Binding-site population

##### Spatial distribution and clustering

A user-specified number of binding sites *N*_*BS*_ are distributed within the nuclear volume following a user-specified cluster-size distribution. For simplicity, the user defines up to five cluster-size bins (binding sites per cluster) and a percentage of total sites in each bin, after which the simulator places clusters whose centres are drawn uniformly at random inside the nucleus. The binding sites for each cluster are drawn uniformly inside a ball of radius *r*_*cluster*_ around the cluster centre. Any sites that would fall outside the nuclear envelope are projected back onto the surface. After the requested clusters are placed, any remaining binding sites are distributed uniformly in the nucleus as singletons.

##### Optional synthetic site array

In addition to the endogenous site population, an optional synthetic cluster (synthetic site array) of *N*_*syn*_ sites can be placed; its centre is drawn uniformly inside the inner 70% of the nuclear radius and its members are drawn uniformly inside a ball of radius *r*_*syn*_ around that centre. The synthetic site clusters are distinctly colored in the ground truth images, and their positions over time are recorded for downstream analysis.

##### Binding-site strengths

Each site is assigned a strength multiplier drawn from a five-bin categorical distribution with user-defined multipliers and bin probabilities. Strengths can be drawn either independently per site or correlated within each cluster (all sites in a cluster receive the same strength). Synthetic sites bypass this distribution and receive the user-set synthetic strength.

##### Site dynamics

Each cluster of binding sites is assigned a diffusion coefficient drawn from a Gaussian with mean and standard deviation specified by the user (clipped at a small positive floor). Cluster centroids then move under fractional Brownian motion (fBm) with a Hurst exponent *H* (the default is *H* = 0.25, showing sub-diffusive Rouse-like polymer motion). *B(t)* is a unique zero-mean Gaussian random variable showing the position of the cluster at time t under fBm. The covariance of the trajectory between time *t* and a future timepoint *s* is given by:

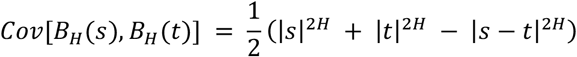

The discrete increments ξ_k_ = *B*(*kΔt*) − *B*((*k*−1)Δ*t*) form the fractional Gaussian noise (fGn), a stationary sequence whose autocovariance at lag *k* (the separation between two increments) is:

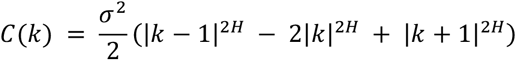

where σ^2^ = 2*D*Δ*t* is the per-step variance and *D* is the cluster diffusion coefficient. The normalized autocovariance γ(*k*) = *C*(*k*)/σ^2^ satisfies γ(0) = 1. For *H* = ½, increments are uncorrelated (standard Brownian motion); *H* < ½ yields sub-diffusive (anti-persistent) increments (default); *H* > ½ yields super-diffusive (persistent) increments. This was implemented via a Davies–Harte construction (SI Methods).

On top of cluster motion, every individual site receives an independent Gaussian “jiggle” around its cluster-translated position at every physics step, to reflect changing conformations of binding sites that are close to each other. Sites are constrained to remain at least one capture radius inside the nuclear surface so that no capture sphere is clipped by the boundary, and jiggled positions are confined to the nucleus.

#### Image formation

##### Voxelization and point-spread function

For each output frame the recorded positions across the exposure window are concatenated and binned onto an isotropic 3D grid with a user-set voxel size, corresponding to the detector pixel size. Each binned position is convolved with an anisotropic 3D Gaussian point-spread function (PSF) with separately specified lateral and axial full-widths at half maximum (*FWHM*_*xy*_, *FWHM*_*z*_). Standard deviations are obtained as 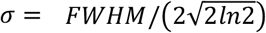. The PSF kernel is truncated at 3*σ* in each axis and normalized to unit mass before deposition such that each molecule represents one intensity unit per frame within the exposure time window (SI Methods). Thus, motion during the exposure window produces blur in the convolved image. Voxels falling outside the nuclear sphere are set to zero.

##### Z-slice projection

The 3D voxel grid is sliced into 2D z-planes spaced at the user-set inter-slice distance or number of slices. Each slice is the single voxel layer at the z-coordinate (no axial integration beyond what the PSF already provides). Slices are saved as TIFF stacks.

#### FRAP Simulations

The molecules in a user-defined spherical region-of-interest are turned to an irreversible dark, “bleached” state. A recovery curve is generated, measuring the intensity of the ROI in the z-plane. The post-FRAP images and intensity values are made available to download.

### Simulations generated for this manuscript

All simulations were generated using the HPC version of SPARK (SI Methods). Bicoid concentrations along the anterior-posterior gradient (20%: 373 nM, 30%: 226 nM, 40%: 137 nM, 50%: 83 nM, and 60%: 51 nM) were estimated from published work^29^. Binding sites were 4392 strong motifs in 670 Bicoid ChIP-seq peaks, assuming 4 chromatid copies, arranged in clusters of 4-24 sites^28^. A small level of cooperativity was included (Hill-like coefficient = 2)^61^. Bicoid clusters in SPARK-generated images were segmented using a previously-published custom segmentation pipeline developed in the lab^8,62^ and were overlaid on the molecule positions to obtain the number of molecules per cluster. Five nuclei were simulated for each concentration and the results were pooled.

Nanog was simulated in 512-cell stage zebrafish nuclei at nuclear concentrations of 5-1500 nM, corresponding to injection of 2-600pg Nanog mRNA. Concentrations were estimated by assuming 10 proteins per mRNA. 9612 binding sites were interpolated from ChIP-seq data at the 256 and 1024-cell stages and two miR-430 arrays of 1200 sites each^31,33–35^. No cooperativity was included.

Sox2 concentrations, kinetics, and the clustering of binding sites in mouse ES cells were obtained from published work^2,37^. Renderings were generated from the positions of only the specifically bound molecules.

Brd4 kinetic parameters were selected by a sweep of exchange rates and on-rate. Free, intermediate and bound fractions were averaged after the simulations reached steady state and compared with experimentally-measured fractions. The point with the smallest maximum error was used for all Brd4 renderings.

Virtual FRAP was performed for the Brd4 simulation. At the bleach frame, every molecule that entered a 1 μm diameter spherical ROI during the exposure was set to a dark state such that it did not contribute any intensity for subsequent frames. ROI intensity was summed over the corresponding disc in the focal plane of the convolved images at the subsequent timepoints and was normalized to the pre-bleach intensity at t = −4s, to match the experimental data^10^. Many examples of cluster fusion were seen across various simulations - the displayed example is for the simulated BRD4 protein. After applying a median filter of 8px and subtracting the result from the original image, clusters were outlined in FIJI on a single z-slice and integrated intensity was measured per frame.

For the binding site simulations (Fig. 4), each simulation contained a single static cluster of synthetic binding sites and no endogenous sites. Cluster occupancy was defined as the number of occupied sites per frame, and turnover is defined as the number of distinct molecules bound to a cluster during a particular length of time.

Sample parameter sets for each displayed example can be found and reused from the dropdown menu of the GUI, or in the “interactive_presets.csv” present in the zipped folder. Full parameter sweep files are given with the code.

### Imaging

#### Fly lines used

The fly line used for Bicoid imaging was *yw*; *his2av-mrfp1; BcdE1, egfp-bcd*.

#### Embryo preparation for live imaging

Drosophila embryos were manually dechorionated by gently rolling them on double-sided tape using a blunt needle. Dechorionated embryos were arranged on an agar pad and transferred onto an adhesive heptane-acrylic spot on a 25mm glass coverslip (lattice light-sheet) or the hydrophobic side of a semi-permeable membrane in a 3D printed sample holder (benchtop confocal).

#### Lattice light-sheet imaging

Lattice light-sheet volumetric imaging of Bicoid was performed on a Multimodal Optical Scope with Adaptive Imaging Correction (MOSAIC)^63^. The 488 nm excitation laser was used at 10% (0.552 mW at the back focal plane of the excitation objective) to excite eGFP. A volume of 18.9 µm sampling z-planes spaced 0.3 µm apart was acquired at various positions along the anterior-posterior axis of the embryo, with 30 ms exposure time.

#### Spinning disk confocal imaging

Confocal imaging was performed on an Andor BC43 Spinning Disk Confocal Microscope with a 20x/0.8 NA objective to obtain a full-embryo image. The 488 nm excitation laser was used at 15% (16.9 mW) to excite eGFP. A volume of 34 µm sampling z-planes spaced 0.3 µm apart was acquired, with 100 ms exposure time.

### Image Analysis

Lattice light-sheet images were pre-processed using GPU-accelerated 3D deconvolution via CUDA^64^. Deconvolution was done using a Richardson-Lucy based algorithm with 5 iterations, utilizing point spread functions (PSFs) captured from bead images acquired on the day of imaging to ensure accurate correction.

All images were processed, scaled and visualized in FIJI, Napari, or Imaris. Displayed kymographs were constructed by taking a 600 nm maximum projection across z-slices, generating a kymograph for each column in the red rectangle, and max-projecting the 5 resulting kymographs.

Data from other papers (Fig. 2, 3) was directly approximated from figures using PlotDigitizer^65^.

### Software Development and AI Assistance

SPARK was implemented in Python by the authors with the assistance of a large language model (Anthropic Claude Opus, online and through Claude Code). Claude was used for code generation, GUI creation, editing, and for the suggestion and implementation of algorithms such as the Davies–Harte construction for fractional Brownian motion. All AI-assisted code and text were reviewed, tested, and validated by the authors, who take full responsibility for the correctness of the implementation.

## ACKNOWLEDGEMENTS

We thank all members of the Mir lab for insightful discussions, rigorous testing of the software, and critical feedback on initial versions of the manuscript. We especially thank Dr. Apratim Mukherjee for construction of the lab’s lattice light-sheet microscope and Dr. Samantha Fallacaro for developing the cluster segmentation pipeline used in this work. We thank Dr. Mariko L. Bennett for access to their lab’s Andor BC43 Spinning Disk Confocal Microscope.

## AUTHOR CONTRIBUTIONS

M.M. conceived and supervised the study. M.K. developed software, performed research and imaging. Both authors interpreted data, prepared figures, and wrote the manuscript.

## FUNDING

The work was supported by National Institutes of Health grant DP2HD108775 (M.M.) and Howard Hughes Medical Institute, Freeman Hrabowski Scholars Program (M.M).

## DATA AND CODE AVAILABILITY

### Data Availability

All simulated data can be reproduced using the provided code.

### Code Availability

Developed code is available on Zenodo: https://doi.org/10.5281/zenodo.23964546

## Notes

https://doi.org/10.5281/zenodo.23964546

### Competing Interest Statement

The authors have declared no competing interest.

### Summary of Updates

All sections, Figs. 1-2, and Fig. S3 updated. Figs. 3-4 and Movie S3 added.

## REFERENCES

1. Meeussen, J. V. W. & Lenstra, T. L. Time will tell: comparing timescales to gain insight into transcriptional bursting. Trends Genet 40, 160–174 (2024).

2. Chen, J. et al. Single-molecule dynamics of enhanceosome assembly in embryonic stem cells. Cell 156, 1274–1285 (2014).

3. Mir, M. et al. Dense Bicoid hubs accentuate binding along the morphogen gradient. Genes Dev 31, 1784–1794 (2017).

4. Mir, M. et al. Dynamic multifactor hubs interact transiently with sites of active transcription in embryos. Elife 7, (2018).

5. Wagh, K., Stavreva, D. A., Upadhyaya, A. & Hager, G. L. Transcription Factor Dynamics: One Molecule at a Time. Annu Rev Cell Dev Biol 39, 277–305 (2023).

6. Fallacaro, S. et al. Enhancer binding kinetics explain transcription factor hub formation. Sci Adv 12, eaeh1086 (2026).

7. Kawasaki, K. & Fukaya, T. Functional coordination between transcription factor clustering and gene activity. Mol Cell 83, 1605–1622.e9 (2023).

8. Mukherjee, A. et al. A single cluster of RNA Polymerase II molecules is stably associated with active genes. Nat Commun 17, (2026).

9. Cho, W.-K. et al. Mediator and RNA polymerase II clusters associate in transcription-dependent condensates. Science 361, 412–415 (2018).

10. Sabari, B. R. et al. Coactivator condensation at super-enhancers links phase separation and gene control. Science 361, (2018).

11. Chong, S. et al. Imaging dynamic and selective low-complexity domain interactions that control gene transcription. Science 361, (2018).

12. Pei, G., Lyons, H., Li, P. & Sabari, B. R. Transcription regulation by biomolecular condensates. Nat Rev Mol Cell Biol 26, 213–236 (2025).

13. Du, M. et al. Direct observation of a condensate effect on super-enhancer controlled gene bursting. Cell 187, 2595–2598 (2024).

14. Alberti, S. et al. Current practices in the study of biomolecular condensates: a community comment. Nat Commun 16, 7730 (2025).

15. Navalkar, A., Eppert, M., Sabari, B. R. & Mittag, T. Density transitions in the regulation of transcription. Mol Cell 86, 567–584 (2026).

16. Boija, A. et al. Transcription Factors Activate Genes through the Phase-Separation Capacity of Their Activation Domains. Cell 175, 1842–1855.e16 (2018).

17. Zamudio, A. V. et al. Mediator Condensates Localize Signaling Factors to Key Cell Identity Genes. Mol Cell 76, 753–766.e6 (2019).

18. Klosin, A. et al. Phase separation provides a mechanism to reduce noise in cells. Science 367, 464–468 (2020).

19. Brangwynne, C. P. et al. Germline P granules are liquid droplets that localize by controlled dissolution/condensation. Science 324, 1729–1732 (2009).

20. McSwiggen, D. T., Mir, M., Darzacq, X. & Tjian, R. Evaluating phase separation in live cells: diagnosis, caveats, and functional consequences. Genes Dev 33, 1619–1634 (2019).

21. Alberti, S., Gladfelter, A. & Mittag, T. Considerations and Challenges in Studying Liquid-Liquid Phase Separation and Biomolecular Condensates. Cell 176, 419–434 (2019).

22. McSwiggen, D. T. et al. Evidence for DNA-mediated nuclear compartmentalization distinct from phase separation. Elife 8, (2019).

23. Gaskill, M. M. et al. Localization of the Drosophila pioneer factor GAF to subnuclear foci is driven by DNA binding and required to silence satellite repeat expression. Dev Cell 58, 1610–1624.e8 (2023).

24. Donovan, B. T. et al. Live-cell imaging reveals the interplay between transcription factors, nucleosomes, and bursting. EMBO J 38, (2019).

25. Alamos, S. et al. Minimal synthetic enhancers reveal control of the probability of transcriptional engagement and its timing by a morphogen gradient. Cell Syst 14, 220–236.e3 (2023).

26. Cadar, E., Birnie, A., Marsman, R., Grond, R. & Bothma, J. P. Transcription factor clusters enhance gene expression in vivo without modulating bursting dynamics. Mol Cell 86, 4234–4249.e9 (2026).

27. Gregor, T., Tank, D. W., Wieschaus, E. F. & Bialek, W. Probing the limits to positional information. Cell 130, 153–164 (2007).

28. Hannon, C. E., Blythe, S. A. & Wieschaus, E. F. Concentration dependent chromatin states induced by the bicoid morphogen gradient. Elife 6, (2017).

29. Munshi, R., Ling, J., Ryabichko, S., Wieschaus, E. F. & Gregor, T. Transcription factor clusters as information transfer agents. Sci Adv 11, eadp3251 (2025).

30. Dongre, S. & Vastenhouw, N. L. Specificity in clustering of gene-specific transcription factors is encoded in the genome. Nucleic Acids Res 53, (2025).

31. Chabot, N. M. et al. Local DNA compaction creates TF-DNA clusters that enable transcription. bioRxiv (2024) doi:10.1101/2024.07.25.605169.

32. Reisser, M. et al. Single-molecule imaging correlates decreasing nuclear volume with increasing TF-chromatin associations during zebrafish development. Nat Commun 9, 5218 (2018).

33. Wang, X. et al. Antibody-free profiling of transcription factor occupancy during early embryogenesis by FitCUT&RUN. Genome Res 32, 378–388 (2022).

34. Xu, C. et al. Nanog-like regulates endoderm formation through the Mxtx2-Nodal pathway. Dev Cell 22, 625–638 (2012).

35. Hadzhiev, Y. et al. The miR-430 locus with extreme promoter density forms a transcription body during the minor wave of zygotic genome activation. Dev Cell 58, 155–170.e8 (2023).

36. Okamoto, K. et al. Single-molecule tracking of Nanog and Oct4 in living mouse embryonic stem cells uncovers a feedback mechanism of pluripotency maintenance. EMBO J 42, e112305 (2023).

37. Liu, Z. et al. 3D imaging of Sox2 enhancer clusters in embryonic stem cells. Elife 3, e04236 (2014).

38. Dey, A., Chitsaz, F., Abbasi, A., Misteli, T. & Ozato, K. The double bromodomain protein Brd4 binds to acetylated chromatin during interphase and mitosis. Proc Natl Acad Sci U S A 100, 8758–8763 (2003).

39. Cho, C.-Y. & O’Farrell, P. H. Stepwise modifications of transcriptional hubs link pioneer factor activity to a burst of transcription. Nat Commun 14, 4848 (2023).

40. Strom, A. R. et al. Interplay of condensation and chromatin binding underlies BRD4 targeting. Mol Biol Cell 35, ar88 (2024).

41. Xie, L. et al. BRD2 compartmentalizes the accessible genome. Nat Genet 54, 481–491 (2022).

42. Driever, W., Thoma, G. & Nüsslein-Volhard, C. Determination of spatial domains of zygotic gene expression in the Drosophila embryo by the affinity of binding sites for the bicoid morphogen. Nature 340, 363–367 (1989).

43. Mullins, M. C. et al. Genes establishing dorsoventral pattern formation in the zebrafish embryo: the ventral specifying genes. Development 123, 81–93 (1996).

44. Jiang, J. & Levine, M. Binding affinities and cooperative interactions with bHLH activators delimit threshold responses to the dorsal gradient morphogen. Cell 72, 741–752 (1993).

45. Farley, E. K., Olson, K. M., Zhang, W., Rokhsar, D. S. & Levine, M. S. Syntax compensates for poor binding sites to encode tissue specificity of developmental enhancers. Proc Natl Acad Sci U S A 113, 6508–6513 (2016).

46. Lim, F. et al. Affinity-optimizing enhancer variants disrupt development. Nature 626, 151–159 (2024).

47. Crocker, J. et al. Low affinity binding site clusters confer hox specificity and regulatory robustness. Cell 160, 191–203 (2015).

48. Crocker, J., Noon, E. P.-B. & Stern, D. L. The Soft Touch: Low-Affinity Transcription Factor Binding Sites in Development and Evolution. Curr Top Dev Biol 117, 455–469 (2016).

49. Zhang, C. et al. A clustering property of highly-degenerate transcription factor binding sites in the mammalian genome. Nucleic Acids Res 34, 2238–2246 (2006).

50. Leith, J. S. et al. Sequence-dependent sliding kinetics of p53. Proc Natl Acad Sci U S A 109, 16552–16557 (2012).

51. Elf, J.Li, G.-W. & Xie, X. S. Probing transcription factor dynamics at the single-molecule level in a living cell. Science 316, 1191–1194 (2007).

52. Wei, Z. et al. Nucleosomes and IDRs suppress promiscuous GCN4 binding on minichromosomes. Nat Struct Mol Biol 33, 1332–1340 (2026).

53. Garcia, D. A. et al. An intrinsically disordered region-mediated confinement state contributes to the dynamics and function of transcription factors. Mol Cell 81, 1484–1498.e6 (2021).

54. Peer, S. & Levy, Y. Disordered Tails Shape DNA Specificity of Myc:Max via Transient Competition. J Mol Biol 438, 169891 (2026).

55. Fallacaro, S. et al. A fine kinetic balance of interactions directs transcription factor hubs to genes. bioRxiv (2024) doi:10.1101/2024.04.16.589811.

56. Brodsky, S. et al. Intrinsically Disordered Regions Direct Transcription Factor In Vivo Binding Specificity. Mol Cell 79, 459–471.e4 (2020).

57. Abidi, A. A. et al. Unstructured transcription factor interactions enable emergent specificity. Science 392, eaeb6487 (2026).

58. Wang, Z. et al. Mesoscale chromatin confinement facilitates target search of pioneer transcription factors in live cells. Nat Struct Mol Biol 32, 125–136 (2025).

59. Strugo, N. et al. Intrinsically disordered regions facilitate target search to drive promoter selectivity by a yeast transcription factor. Nat Commun 17, 523 (2025).

60. Sakong, S., Fierz, B. & Suter, D. M. Electrostatic properties of disordered regions control transcription factor search and pioneer activity. Nat Commun 17, (2026).

61. Burz, D. S., Rivera-Pomar, R., Jäckle, H. & Hanes, S. D. Cooperative DNA-binding by Bicoid provides a mechanism for threshold-dependent gene activation in the Drosophila embryo. EMBO J 17, 5998–6009 (1998).

62. Fallacaro, S. et al. Enhancer binding kinetics explain transcription factor hub formation. bioRxiv (2026) doi:10.1101/2025.04.07.647578.

63. Fu, T.-M. et al. A multimodal adaptive optical microscope for in vivo imaging from molecules to organisms. Nat Methods https://doi.org/10.1038/s41592-026-03066-1 (2026) doi:10.1038/s41592-026-03066-1.

64. GitHub-scopetools/cudadecon: GPU accelerated 3D image deconvolution using CUDA. GitHub https://github.com/scopetools/cudadecon.

65. PlotDigitizer, 3.1.6, 2026. https://plotdigitizer.com.

