## Supplemental Information for "Molecular diffusion and binding kinetics are sufficient to explain condensate-like nuclear clustering"

### SI Methods

Davies–Harte construction for fBm.

fGn increments for the whole run are generated simultaneously for all clusters and all three spatial axes via the circulant embedding of the autocovariance matrix:

(1) Form the circulant embedding vector of length $N = 2n$:

$$r_j = \gamma(j),\ 0 \le j \le n;\quad r_j = \gamma(2n - j),\ n < j < 2n$$

(2) Compute eigenvalues $\lambda_j = \mathrm{Re}[\mathrm{DFT}(r)]_j$; clip any negative values to zero.

(3) Construct random vector $W$ of length $N$ from independent standard normals $Z$:

$$W_0 = \sqrt{\frac{\lambda_0}{N}}Z_0,\quad W_n = \sqrt{\frac{\lambda_n}{N}}Z_n$$

$$W_j = \sqrt{\frac{\lambda_j}{2N}}\left(Z^{(1)} + iZ^{(2)}\right),\ j = 1, \ldots, n-1;\quad W_{N-j} = \overline{W_j}$$

(4) The first $n$ elements of $\mathrm{Re}[\mathrm{DFT}(W)]$ are exact unit-variance fGn samples $\{y_k\}$.

(5) Scale to physical units: $\Delta x_k = \sigma\, y_k = \sqrt{(2D\Delta t)}\, y_k$.

Intensity normalization

Two mutually exclusive normalization modes are supported:

1. **Auto-normalize.** All rows are simulated and a single global $v_{max}$ equal to max voxel intensity across every frame is then used to normalize every output. This guarantees intensity comparability across rows. A noise floor as a percentage of the $v_{max}$ can be added. For the batch mode, the normalization is across all conditions batched. A headroom factor can be added for visualization.
2. **Arbitrary-units (AU) calibration.** When AU mode is enabled, the 3D grid is rescaled by a single factor per simulation, chosen so that the mean intensity inside the nuclear mask in the final frame matches an expected signal $S = \left(S_{ref} \,.\, c/c_{ref}\right)$ derived from a user-supplied reference concentration-signal pair. A constant background and Gaussian noise are added inside the nucleus. Output is clipped to the chosen bit depth. AU calibration is set per row; rows with it enabled override the global brightness mode.

Output products

Three image products may be saved per frame (as TIFFs):

- **Convolved.** The PSF-convolved z-slice stack described above, downloaded as 8- or 16- bit. This is the primary simulated microscopy output.
- **Ground truth.** A scatter plot of true molecule centroids (coloured by state) overlaid with binding-site positions (coloured by strength, with synthetic sites highlighted), rendered at a user-specified upscale factor for visual inspection.
- **Molecule count map.** A z-slice stack containing, per voxel, the raw count of molecules present there during the exposure window — the same geometry as the convolved stack but without PSF convolution. Three count modes are provided: *full exposure* (every sub-step during the exposure

window contributes one count per molecule, so a molecule present for $N$ sub-steps is counted $N$ times), *snapshot* (only the final sub-step is counted, giving instantaneous voxel occupancy) and centroid (an average of the sub-steps is shown). Each molecule is assigned to the nearest z-slice and counted if it lies within $\pm$(slice-spacing)/2.

A CSV containing the positions and state of each molecule in each frame is also available to download. When a synthetic site array is present, a centroid CSV is also written, giving the centre-of-mass of the synthetic cluster. Finally, a .pkl file that saves the raw simulation can be downloaded, allowing users to upload it to perform FRAP at a later time.

Reproducibility

To support exact replay and to support comparing nominally identical conditions across rendering parameters, the simulator has an integer-seed mechanism that makes every random draw — site positions, strengths, per-cluster diffusion coefficients, fBm increments and jiggle, initial molecule placements, state transitions, binding dice, and residence-time draws — deterministic.

*Interactive mode*
A "Use random seed (reproducible)" checkbox in the General Parameters panel exposes an integer seed input. When checked, the engine calls numpy.random.seed(seed) once at entry, before any random draw. The user can choose whether to freeze the trajectories (visualization parameters can be changed) or only the binding site characteristics (molecule kinetic parameters can also be changed).

*Batch mode*
Per-row reproducibility is controlled by three optional CSV columns:

- **sim_seed** — integer random number generator seed for the row.
- **sim_group** — string label tying together rows that should share *everything* (sites, motion, binding events). Only rendering parameters (PSF FWHMs, voxel size, slice spacing, bit depth, brightness, AU parameters) may differ within a sim_group.
- **site_group** — string label tying together rows that should share *only* the site layout (positions, cluster IDs, strengths, per-site diffusion coefficients). Motion physics — concentration, kinetics, diffusion coefficients — may vary within a site_group. If site_group is blank, it is auto-derived from sim_group.

The per-row seed is resolved as: (1) the row's sim_seed cell, if a usable integer; otherwise (2) base_seed + row_index, where base_seed is the UI-supplied base integer (mirroring the SLURM --base-seed convention); otherwise (3) operating-system entropy. In the HPC implementation, the sim_seed cell is used only for rows that belong to a sim_group or site_group. Ungrouped rows use base_seed + row_index or if no base_seed is supplied, operating-system entropy. Because every random draw inside the engine is reached deterministically from a single global seed, two rows that share a sim_group and seed produce identical output, and two rows that share a site_group and seed produce identical sites with independent molecule motion if concentrations/kinetic parameters differ — the same operational guarantee as the HPC implementation, achieved without an explicit cache. The seed used for every row is logged to seeds_used.csv inside the output ZIP.

Batch processing

A batch run is driven by a CSV in which each row is a complete parameter set. A pre-filled template is generated by the application from the same defaults used by the interactive mode, with the optional reproducibility columns (sim_seed, sim_group, site_group) appended at the end. Rows are processed sequentially and outputs are bundled into a single downloadable ZIP with a top-level subfolder per row. When auto-normalize brightness mode is selected, the simulator runs in two phases: all rows are simulated and gridded in phase 1 to establish the global maximum intensity; phase 2 then renders all rows against that single $v_{max}$ so that intensities are directly comparable across the entire batch.

<u>HPC implementation</u>

The same physics is also implemented as a standalone command-line tool for parallel processing on an HPC. There are two operational differences:

1. The HPC tool uses one SLURM array task per CSV row, so all rows run in parallel under independent processes (unless there are sim or site-group leaders present, which run first). The application processes rows sequentially in the browser session. Because every row is fully determined by its own seed and parameters, the two execution orders produce equivalent output.
2. The HPC tool implements an explicit ground-truth cache: the first row in each sim_group writes its full simulation to disk and the subsequent rows in the same group skip simulation entirely and re-render from the cache. The application does not implement this cache: every row is re-simulated. However, because seeded rows are deterministic, the *output* of sim_group and site_group membership is preserved — rows that share a sim_group and seed produce identical output in the application; rows that share a site_group and seed produce identical sites with independent motion. The application is thus slower than the HPC tool on grouped batches but produces matching results.

### SI Figures

A.

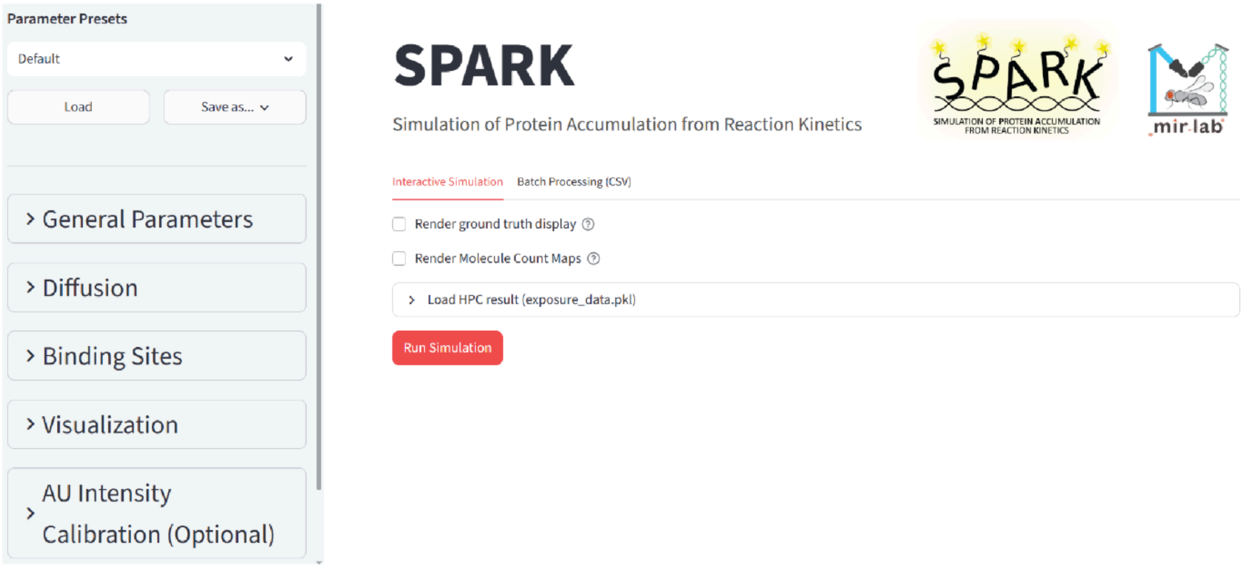


B.

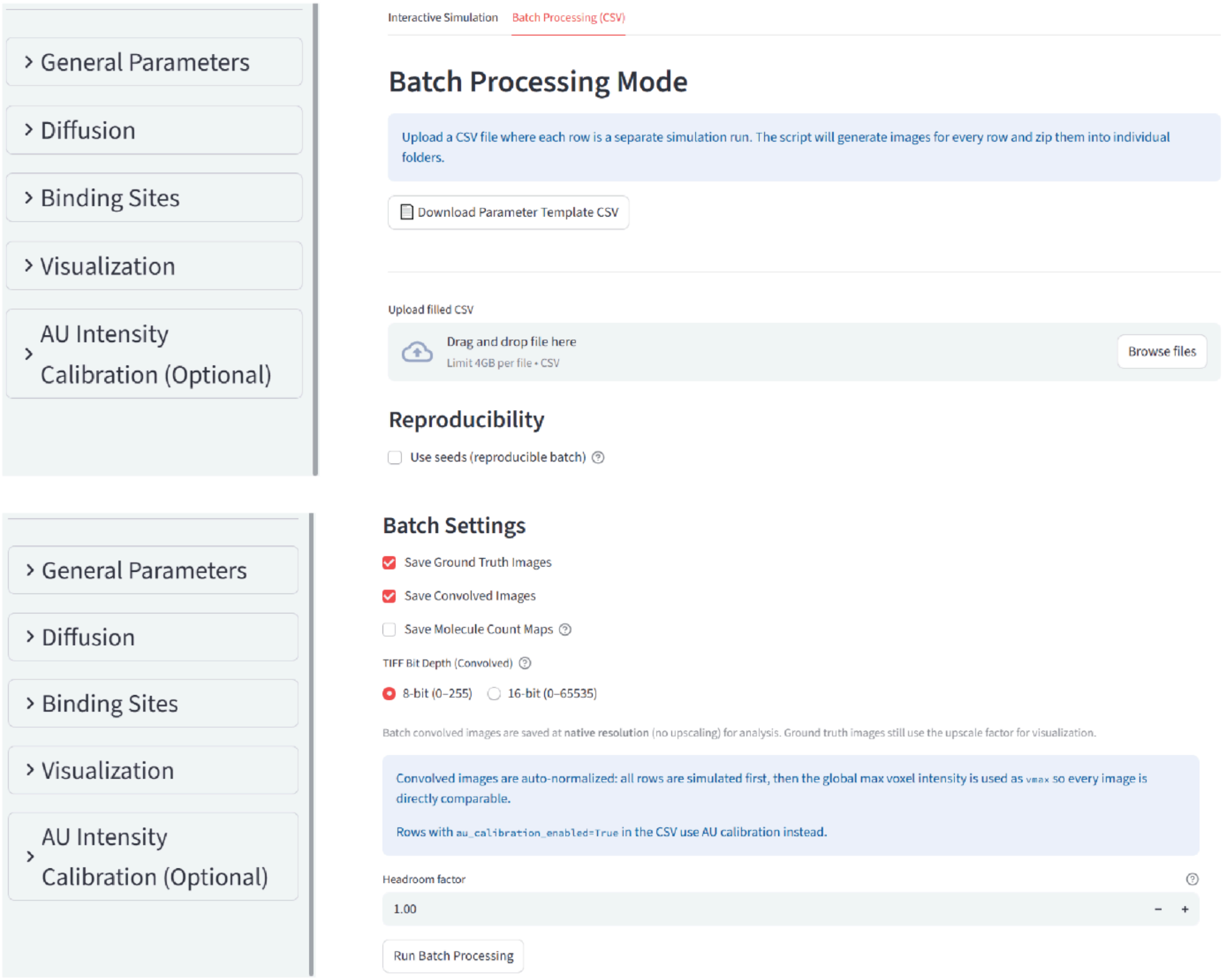


**Fig. S1:** Screenshots of GUI in either **(A)** Interactive or **(B)** Batch Processing mode.

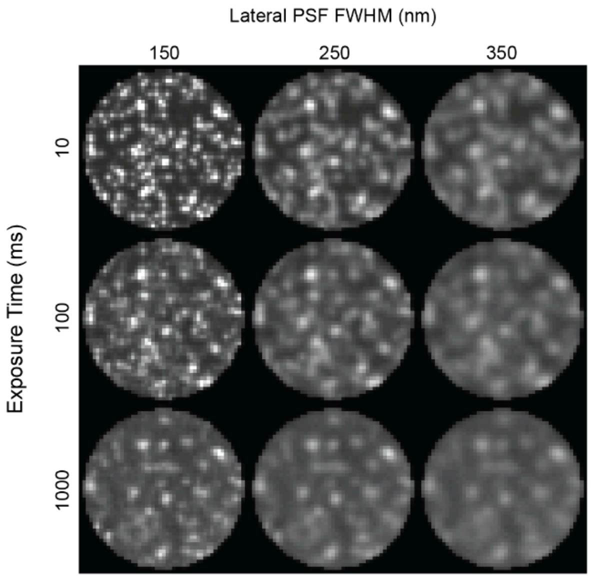


**Fig. S2:** Single timepoint, single z-slice images of nuclei with identical molecular kinetics, showing the effect of modulating either the lateral point spread function or the exposure time. Nucleus diameter is 5 µm.

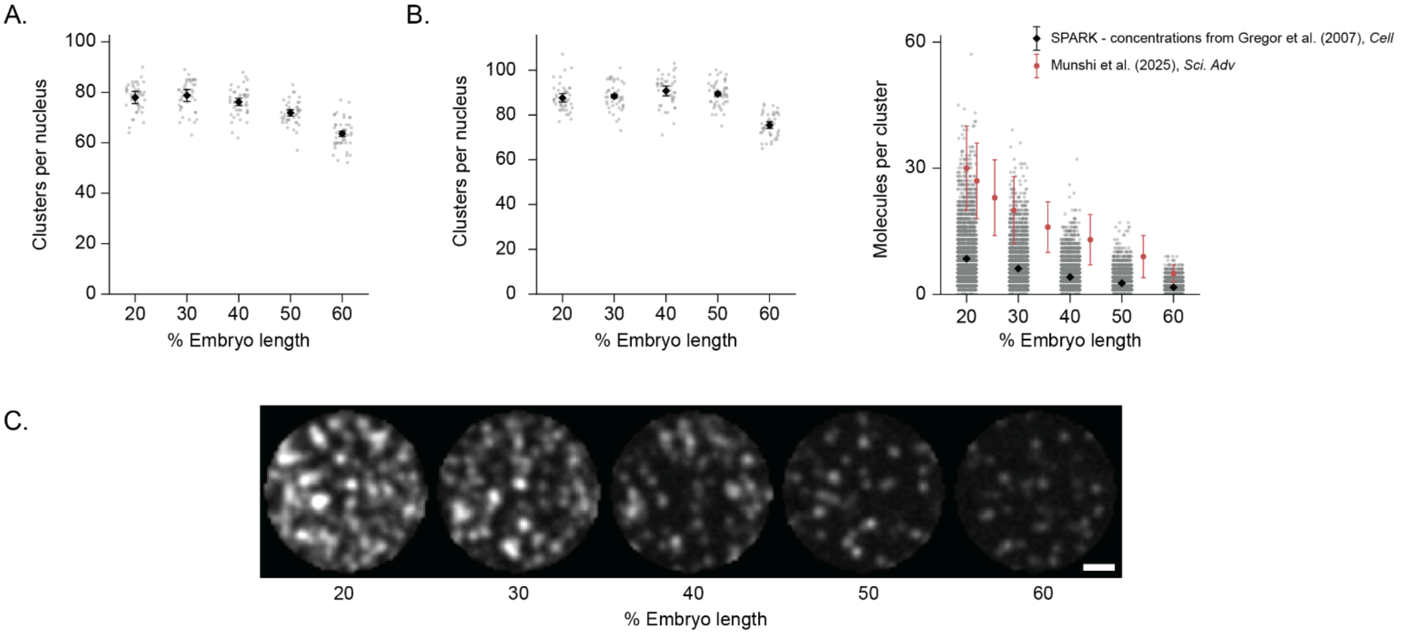


**Fig. S3: (A)** Number of Bicoid clusters per nucleus, corresponding to data in Fig. 2E-G. **(B)** Number of Bicoid clusters per nucleus (left) and molecules per cluster (right) for nuclei with concentrations estimated from Gregor et al.[27]. **(C)** Sample images of Bicoid nuclei using the concentration estimates in (B). Scale bar is 1 μm.

#### Movies

Movie S1: 3D rendering of specifically-bound Sox2 molecules in mouse ES cell nuclei with non-clustered (left) and clustered (right) binding sites.

Movie S2: Simulated FRAP from Fig. 3E. The red circle indicates the bleached area. Scale bar is 1 μm. The duration of the movie is 30 sec. The movie was rendered at 1 fps.

Movie S3: Example fusion event from Fig. 3G. The red circle indicates the fusing clusters. Scale bar is 1 μm. The duration of the movie is 18 sec. The movie was rendered at 1 fps.
